# PeakVI: A Deep Generative Model for Single Cell Chromatin Accessibility Analysis

**DOI:** 10.1101/2021.04.29.442020

**Authors:** Tal Ashuach, Daniel A. Reidenbach, Adam Gayoso, Nir Yosef

## Abstract

Single-cell ATAC sequencing (scATAC-seq) is a powerful and increasingly popular technique to explore the regulatory landscape of heterogeneous cellular populations. However, the high noise levels, degree of sparsity, and scale of the generated data make its analysis challenging. Here we present PeakVI, a probabilistic framework that leverages deep neural networks to analyze scATAC-seq data. PeakVI fits an informative latent space that preserves biological heterogeneity while correcting batch effects and accounting for technical effects such as library size and region-specific biases. Additionally, PeakVI provides a technique for identifying differential accessibility at a single region resolution, which can be used for cell-type annotation as well as identification of key cis-regulatory elements. We use public datasets to demonstrate that PeakVI is scalable, stable, robust to low-quality data, and outperforms current analysis methods on a range of critical analysis tasks. PeakVI is publicly available and implemented in the scvi-tools framework: https://docs.scvi-tools.org/.

## Introduction

Regulatory elements in the genome tend to reside in regions of open chromatin, making the landscape of chromatin accessibility a valuable target of study. Several molecular assays have been developed to support this effort [1, 2, 3], among them ATAC-seq [4], in which accessible regions are fragmented, and the corresponding DNA fragments are sequenced and mapped back to the reference genome, accumulating in areas of open chromatin. Recent advances in sequencing technologies enable performing this assay in single cells [5], thereby allowing the study of chromatin variability at a single cell resolution. Application of Single-cell ATAC-seq (scATAC-seq) has led to promising results in discerning sources of variation, beyond those observed at the transcriptional level [6, 7] and allowed for high resolution characterization of the regulation of in continuous processes, e.g., in immunity [6].

Despite the potential of scATAC-seq, analyzing the resulting data remains challenging. scATAC-seq assays have generally limited sensitivity, detecting 5-15% of accessible regions [7], a common issue for single cell genomics. Additionally, the coverage of this data is limited a-priori since each genomic region has at most two copies in a single cell. Finally, scATAC-seq is extremely high-dimensional, often consisting of hundreds of thousands of genomic regions. These challenges require specialized processing and analysis methods that are designed to account for the specific properties of scATAC-seq data.

One common task for analyzing scATAC-seq is dimensionality reduction: transforming the data to a low-dimensional space that preserves the meaningful information in the original data. This step is crucial to make some downstream analyses, such as clustering and visualization, less noisy, more stable, and computationally tractable. Existing methods use various approaches to achieve this task. Some use methods developed for natural language processing (e.g., latent Dirichlet allocation used by cisTopic [8] and latent semantic analysis (LSA) used by ArchR [9]) that inherently handle sparse high-dimensional data, but do not inherently account for confounding factors that do not have an analog in textual language, such as batch effects. Other methods reduce dimensionality by first aggregating individual regions in the scATAC-seq data to easily interpretable features, such as binding motif scores in the case of chromVAR [10] or gene activity scores in the case of Cicero [11], which makes the data easier to analyze but masks the fine-grain single-region resolution provided by scATAC-seq. These methods have been demonstrated to be under-powered in capturing the true heterogeneity in the original data [12]. Finally, recent methods use deep generative models (e.g., SCALE [13]), but do not account for technical factors, and suffer from model over-fitting due to the dimensionality of the data in contrast with the limited number of samples.

Another common task is differential accessibility analysis. The ability to identify chromatin regions that are preferentially accessible in one population compared with another is foundational to characterizing the chromatin remodelling between cellular identities and states. However, specialized methods to perform this task in the context of scATAC-seq data have not yet been developed. Methods that rely on aggregation of individual regions, like chromVAR and Cicero, perform differential analyses in the aggregated space, thereby losing the single-region resolution. Other methods use linear models developed for RNA-seq data [14] or standard statistical tests [9]. These approaches often suffer from numerical instability due to the sparsity of the data, and statistically overpowered due to the large sample size.

Some recent processing pipelines, like SnapATAC [14] and ArchR [9], offer comprehensive end-to-end analysis pipelines that resolve many issues with processing scATAC-seq data, such as sensitive peak-calling, promoter-enhancer association, and doublet detection. However, these pipelines rely on sub-optimal algorithms for the fundamental tasks mentioned above, and can therefore be improved with better performing methods for those tasks.

Here we present PeakVI, a deep generative model that learns a probabilistic low-dimensional representation of single cells from their chromatin accessibility landscape. PeakVI accounts for technical biases in the data stemming from batch effects, variation in sequence coverage, and bias due to the width of DNA regions, and creates a representation of the data that minimizes these effects. The representation is provided at two levels. One part of the model infers a representation for each cell in a latent low-dimensional space. This latent representation and the space it is embedded in can be used directly for downstream analyses: integration of data sets, identification of cellular sub-populations, and visualization. A second part of the model provides a corrected, probabilistic representation of the raw data. This high dimensional representation enables statistically robust inference of single region-level differential accessibility and cell state annotation. We demonstrate PeakVI’s performance on published data and benchmark it against state-of-the-art published methods on a range of analysis tasks. We show that PeakVI is a powerful addition to the arsenal of scATAC-seq methods and provides capabilities that can help unlock the full potential of scATAC-seq data analysis. PeakVI is publicly available as part of the scvi-tools [15] suite of deep generative models for single cell genomics.

## Results

### 1 PeakVI Model

PeakVI leverages variational inference with deep neural networks to model scATAC-seq data. For each cell, PeakVI estimates the probability of each chromatin region being accessible, as well as technical factors that affect the probability of an accessible region being observed. The standard output of most scATAC-seq preprocessing pipelines (including those employed here; see Methods) is a table of *N* cells and *K* genomic regions. The regions typically correspond to DNA segments with enriched accessibility that are inferred through peak-calling over cell aggregates [16, 14, 9].

The starting point of PeakVI is therefore a *N × K* matrix *X* where *x*_*ij*_ is the number of reads from cell *i* that map to region *j*. While these observations are counts, the underlying biology is mostly binary (a region is either accessible or not). Therefore, PeakVI models the observations as samples from a Bernoulli distribution *P*(*x*_*ij*_ > 0 | *y*_*ij*_, *r*_*j*_, *ℓ*_*i*_), where *y*_*ij*_ is the probability of region *j* being accessible in cell *i*, *r*_*j*_ ∈ [0, 1] is a region-specific scaling factor, and *ℓ*_*i*_ ∈ [0, 1] is a cell-specific scaling factor (Figure 1A). Conceptually, these components are related to the three molecular events that are required for a region to be observed as accessible: (1) the region must be accessible in the cell, which largely depends on the cell state and identity, captured by *y*_*ij*_; (2) the accessible region must be tagmented by the transposase that underlies the ATAC-seq protocol, a process which may be skewed by region-specific factors such as width (in base pairs) and sequence biases, captured by *r*_*j*_; (3) finally, the corresponding fragment must be captured and sequenced, which may also depend on library-specific factors, such as sequencing depth and efficacy of the library preparation, captured by *ℓ*_*i*_.

**Figure 1:**
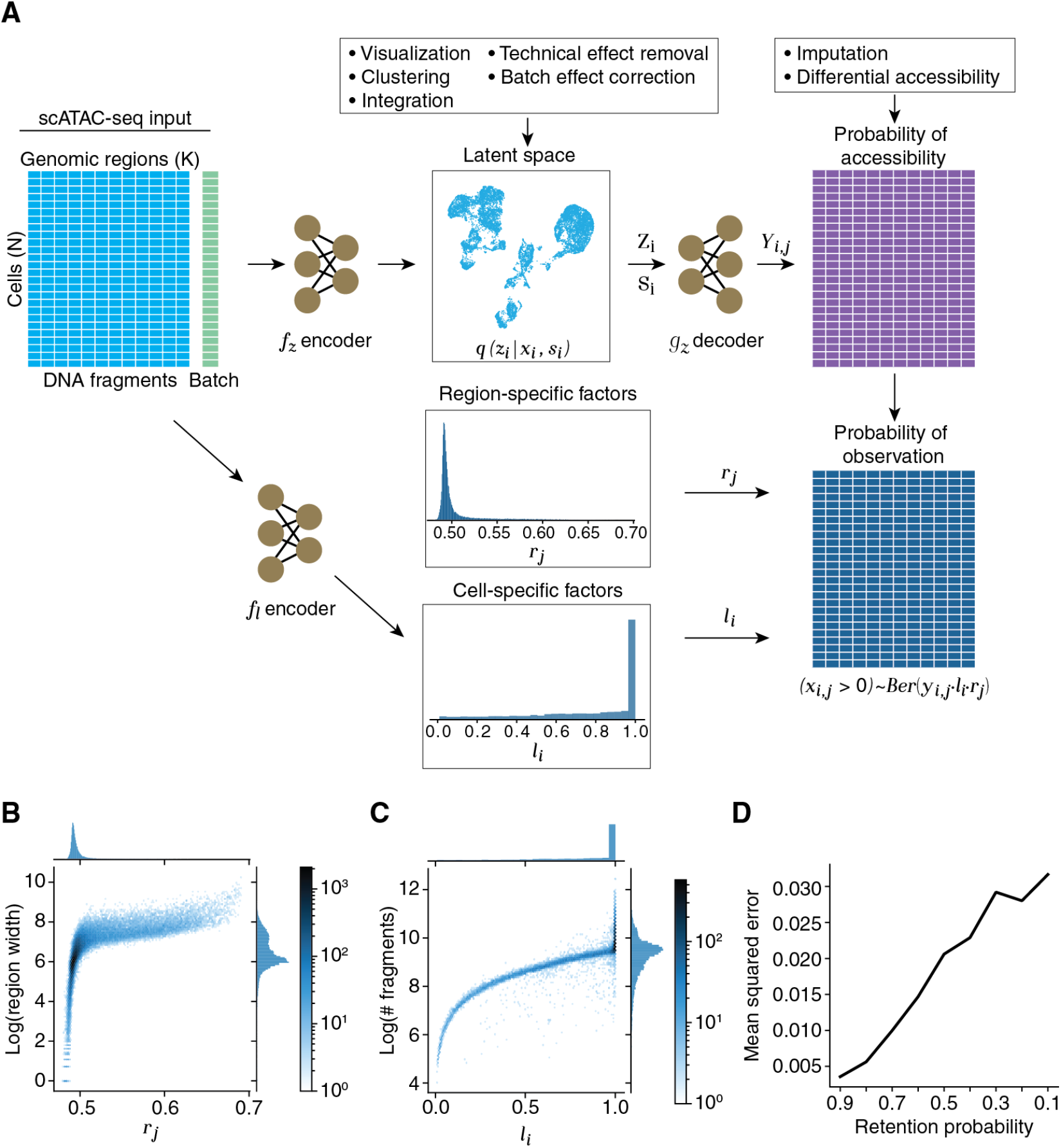
PeakVI Model overview. **(A)** conceptual model illustration. The input region-by-cell count matrix (left) is estimated as the product of region-specific effects (center top), cell-specific effects (center), and accessibility probability estimates (center bottom). The observation probability matrix (right) is used to calculate the likelihood of the data for optimization. **(B)** The region-specific factor *r*_*j*_ is assigned higher values for wider regions, indicating a higher probability of those regions being fragmented. **(C)** The cell-specific factor *ℓ*_*i*_ increases with the number of fragments up to a saturation point. Cells with sufficient fragments are not penalized even if other cells have significantly more fragments. **(D)** Random corruption of the data at increasing rates leads to a small but steady increase in the mean squared error (measured from corrupted indices).

PeakVI uses a variational autoencoder [17] (VAE) and an auxiliary neural network to estimate these factors. The VAE consists of two major components: (1) the encoder network *f*_*z*_ infers the distributional parameters of the *d*-dimensional (for *d ≪ D*) latent representation *z*_*i*_ (also known as the variational posterior) from the observed data: *f*_*z*_(*x*_*i*_ = *q*(*z*_*i*_|*x*_*i*_); (2) the decoder network *g*_*z*_ and the generative model, which takes a sample from the latent representation *z*_*i*_ and the batch annotations *s*_*i*_ and generates an estimate of the probability of each genomic region being accessible in the cell *i*: (*g*_*z*_(*z*_*i*_, *s*_*i*_))_*j*_ = *y*_*ij*_. The cell-specific scaling factor *ℓ*_*i*_ is inferred from the observed data using an additional neural net *f*_*ℓ*_, and the region-specific scaling factor *r*_*j*_ ∈ [0, 1] is optimized directly as a model parameter. Finally, the probability of observing a region in a cell (i.e *p*(*x*_*ij*_ > 0)) are computed as the product of the three probabilities: *p*(*x*_*ij*_ > 0) = *y*_*ij*_ · *ℓ*_*i*_ · *r*_*j*_ (Figure 1A). Formally:

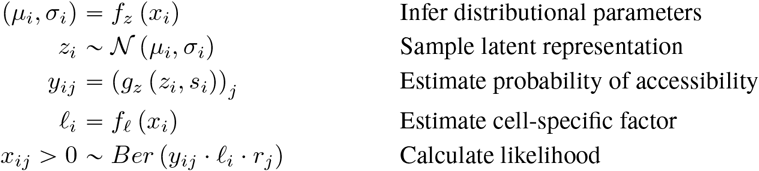

Conditioning on batch annotations, or any other known sources of unwanted variation, encourages the encoder to capture batch-independent biological variation in the latent representation *z*_*i*_, which can then be used for normalized and batch-corrected visualization, clustering, and other downstream analyses. The inferred accessibility probabilities *y*_*ij*_ are an estimate of the true chromatin landscape in each cell, while technical effects that stem from either region-specific biases or cell-specific biases are captured by the *r* and *ℓ* scaling factors, respectively. We can then estimate the probability of observing a region in each cell as the product of these factors *y*_*ij*_ · *ℓ*_*i*_ · *r*_*j*_ and compute the likelihood of the observations. During training, a lower bound of the marginal log likelihood log *p(x*_*ij*_ > 0) is then maximized using auto-encoding variational Bayes [17]. Full model architecture and training parameters are provided in the Methods section.

#### 1.1 Benchmark Datasets

In order to evaluate the performance of PeakVI, we examined both simulated and real datasets. We found, however, that current simulation techniques [12] rely on independent sampling from distributions attained from bulk ATAC-seq data, which creates a highly-sparse covariance structure that does not realistically reflect assayed datasets (Figure S1). Our analysis therefore relies primarily on two publicly available datasets: (1) Hematopoiesis data from Satpathy et al. [6] which consists of bone marrow and blood samples that were flow-sorted for different cell subsets, as well as several batches of unsorted samples that consist of multiple cell types; (2) A dataset released by 10x Genomics of joint RNA-seq and ATAC-seq from single human peripheral blood mono-nuclear cells (PBMCs). The first dataset contains cell type specific labels that provide an established benchmark, as well as multiple batches that allow comparison of batch effect correction. The second dataset provides an orthogonal modality of data that can be used to validate scATAC-based analyses. Finally, the two datasets are generated using different protocols and are processed differently, allowing us to demonstrate the PeakVI’s performance is protocol- and processing-independent.

#### 1.2 PeakVI Captures Nuanced Effects of Technical Confounders

Since the normalization factors included in the PeakVI model, *r* and *ℓ*, are optimized by the training process, we set out to confirm that they converge on values that correspond to the empirical, technical confounders. We used the 10X PBMC data for these analyses. For the region-specific factor *r*, we examined how it corresponds to the width of the genomic region, a known technical confounder. we found that PeakVI assigns the vast majority of regions with a value around 0.5, with higher values indeed being assigned to wider regions, which have a higher probability of being fragmented (Figure 1B). Notably, the overall distribution of this factor only reaches as high as roughly 0.75, well below the max value of 1. This translates to a global penalty imposed on all observations, which implicitly reflects the limited sensitivity of this assay and the resulting abundance of false-negative observations. For the cell-specific factor *ℓ* we examined how it corresponds to the number of reads captured in each cell. We find that the vast majority of cells have *ℓ* ≈ 1, and the dynamic values of *ℓ* indeed correspond to the empirical library size (Figure 1C). The saturation of this factor reflects an important consideration when normalizing library sizes for chromatin profiling: different cell types may have different levels of accessibility (e.g unbalanced chromatin remodeling during differentiation [18]), therefore this factor should not penalize cells states with less accessible chromatin, but rather only weigh down cells in cases where the decrease in fragments is due to technical effects. Overall we see that the normalization factors used by the model have a clear but nuanced correspondence to empirical confounders.

#### 1.3 PeakVI is Robust to Increased Sparsity and Stable Across Hyperparameters

Limited sensitivity, which results in an abundance of missing observations, is a major problem in single-cell assays and particularly scATAC-seq. We therefore examined how PeakVI handles increasing levels of sparsity. We corrupted the 10X PBMC data by randomly replacing non-zero observations with zeros at a range of probabilities (10 90%) and trained PeakVI on each corrupted dataset. We then used PeakVI’s estimates of the probability of accessibility for these corrupted observations and compared the estimates from the models trained on corrupted data, in which these observations were 0, to the original estimates from the model trained on the full data, where these observations where non-zero. We computed the error: 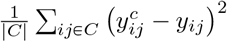, where *C* is the set of corrupted observations, *y*^*c*^ is the probability of accessibility estimated by peakVI when trained on the corrupted data, and *y* is the probability of accessibility estimated from the original, uncorrupted, data. We found that PeakVI produces highly consistent results, even in highly sparse situations: with a mean squared error of 0.06 when 10% of the observations are removed, to 0.17 when 90% of the data is removed (Figure 1D, S2). We also observed that the corrupted estimates are generally lower than the original estimates, consistent with the corruption being one-directional (introducing false negatives, not false positives). These results demonstrate that PeakVI is robust to low-quality and highly sparse data.

Since training PeakVI involves stochastic optimization of a non-convex function, the model can produce different results in different runs. We examined how stable PeakVI is to changes in architecture and training hyperparameters by training PeakVI on a variety of configurations and comparing how the different models perform on held-out data. We varied the number of hidden layers in the neural networks, the size of the mini-batch used in training, the dropout rate, and learning rate, and the weight decay. For each set of hyperparamters, we trained the model 3 times, and measured the likelihood the model achieves on the held-out data in each run. We found that PeakVI is highly stable, and that the default hyperparameters perform well without a need to fine-tune the model for each analysis (Supp. Table 1, Methods).

### 2 PeakVI Learns an Informative Batch-Corrected Latent Representation

PeakVI learns a low-dimensional representation of each cell that preserves biological heterogeneity while reducing noise, technical artifacts, and batch effects. We compared the latent space learned by PeakVI with representations from published methods. We compared to four methods: 1) latent semantic analysis (LSA), a natural language processing (NLP) technique commonly used in scATAC-seq analysis pipelines[14, 9]; 2) cisTopic[8], a method that uses Latent Dirichlet Allocation, commonly used for NLP tasks; 3) SCALE[13], a method that also employs a VAE, and incorporates Gaussian mixture modelling (GMM) to create a clustered latent space; 4) chromVAR[10], an algorithm that aggregates genomic regions by known binding motifs and normalizes these aggregates to motif accessibility scores. The first two methods, LSA and cisTopic, were chosen since a recent benchmark of computational analysis methods for scATAC-seq methods[12] found them to be the best performing methods. SCALE, published after the benchmark, is included in our comparison due to the conceptual similarities with PeakVI. Finally, we included chromVAR since it is commonly used as both a dimensionality reduction method as well as an annotation technique.

First we used the 10X PBMC scATAC-seq data to measure how consistent each latent representation is with the gene expression profiles that are also measured from each cell. We ran all methods on the 10X PBMC data and extracted the latent representation computed by each. We then independently analyzed the paired scRNA-seq data and clustered the cells based on their gene expression profiles (Methods). We then overlaid the scRNA-based cluster labels on the scATAC-based representations (Figure 2A), and measured for each cell the fraction of its chromatin-based K nearest neighbors that are from the same RNA-based cluster for varying values of K (Figure 2B, Methods). We found that PeakVI and cisTopic outperformed all other methods, with PeakVI doing marginally better than cisTopic. We also measured how robust each method is to library size effects, by computing for each latent space the correlation of the latent representation with the empirical library size (log(number of fragments)), using Geary’s C [19] (Figure S3, Methods). We found that LSA and SCALE are especially sensitive to library size effects, while PeakVI and cisTopic are more robust, and chromVAR having a negative association with library size.

**Figure 2:**
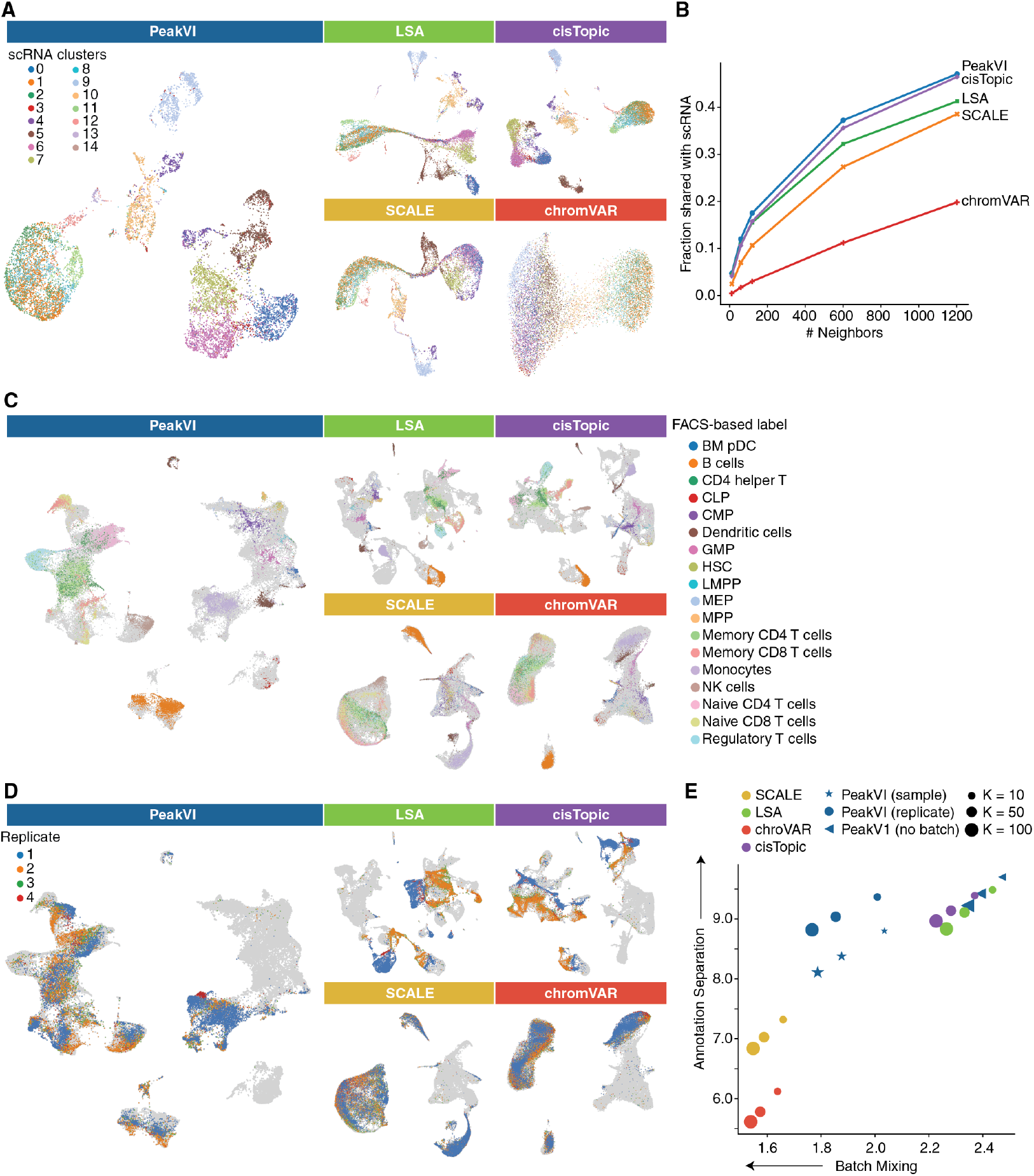
UMAP visualizations of latent representations from PeakVI, LSA, cisTopic, SCALE, and chromVAR. **(A)** The paired scRNA-scATAC sample PBMC dataset from 10X Genomics. Cells are colored based on the scRNA-based clustering, umaps are computed from the scATAC representations. All methods except for chromVAR are comparably consistent with the scRNA data. **(B)** Quantitative consistency of the latent representation with the scRNA data; fraction of the *K* nearest neighbors in the scATAC representation that are also among the *K* nearest neighbors in the scRNA representation, for various values of *K*. PeakVI marginally outperforms cisTopic, followed by LSA, SCALE, and chromVAR. **(C)** Data from Satpathy et al [6]; cells are colored using the FACS-based cell type-specific labels. Cells from unsorted samples or non-specific sorted samples are colored in light gray. PeakVI, LSA, and cisTopic all achieve good separation of cell types. **(D)** Data from Satpathy et al [6]; cells are colored using the unsorted PBMC replicates. Cells from all other samples are colored in light gray. Batch effects are reduced with PeakVI, chromVAR, and SCALE. **(E)** Enrichment of labels among the K-nearest neighbors for each cell; X-axis is the enrichment of batch labels, where lower enrichment indicates better batch mixing. Y-axis is the enrichment of cell type labels, where higher enrichment indicates better separation. PeakVI reaches a better balance of the two tasks.

Next we looked into how each method handles a more complex experimental design, as in the hematopoiesis dataset, which consists of multiple samples in different sizes, some cell type specific and others general. We analyzed the data with all methods, and for PeakVI used three configurations: (1) “no batch”, without any batch annotation; (2) “full batch”, treating each sample as a separate batch; (3) “replicate batch”, treating each replicate from multi-replicate conditions as a separate batch (Methods). These configurations correspond to having no batch correction, strict batch correction, or an intermediate approach, respectively. We examined how well each method preserves biological heterogeneity by measuring how separated the sorted cell populations are, using the cell type-specific fluorescence-based labels (Figure 2C, S4). We also examined how well each method handles batch effects, which none of the examined methods explicitly corrects, by measuring how well-mixed are the four different batches of unsorted PBMC samples (Figure 2D, S4). For both analyses we computed an enrichment score by computing for every cell the number of neighbors out of its *K*-nearest neighbors that share its label, and comparing to the random expectation (Methods), for varying values of *K* (scores in the text are for *K* = 50) (Figure 2E). Ideally, this enrichment score would be high for biological labels and low for batch labels. We find that LSA, cisTopic, and PeakVI with no-batch configuration all achieve high separation (enrichment scores 9.1, 9.13, and 9.42, respectively) but separate the different batches as well (enrichment scores 2.33, 2.28, 2.39 respectively); conversely, chromVAR and SCALE outperform all methods in batch mixing (1.57 and 1.59, respectively), but do worse on cell type separation (5.78 and 7.03, respectively). In contrast, we find that PeakVI with replicate-batch strikes a desirable balance, preserving biological heterogeneity comparably well (enrichment score 9.04) while more effectively mixing the batches (enrichment score 1.85). Finally, PeakVI with full-batch configuration also achieves a good balance (8.37 for cell type separation, 1.88 for batch mixing), but underperforms the replicate-batch configuration on both tasks. Overall these results demonstrate that PeakVI is better able to correct batch effects while preserving biological heterogeneity, reaching an overall better latent representation than all examined methods.

### 3 PeakVI performs differential accessibility analysis at a single-region resolution

Among the main promises of scATAC-seq is the ability to better identify individual genomic elements that help regulate certain biological processes. Achieving this requires the ability to identify individual regions that are differentially accessible between different groups of cells. In practice this task is challenging due to the binary nature of each observation, batch effects, and the high levels of noise and sparsity. Most differential analyses thus choose to aggregate the differential signal across different regions, either by the binding motifs they harbor (i.e the differential analysis chromVAR performs) or by aggregating the surrounding regions to each gene and creating a gene activity score [11]. While these analyses are useful, they do not enable identification of individual regions, thereby not fully unlocking the promise of scATAC-seq data.

PeakVI addresses this problem by leveraging the probabilistic nature of the latent space to produce denoised and normalized estimates of accessibility, which enable a robust and accurate estimate of differential accessibility at a single-region resolution. Briefly, given a population of cells *C* and a region *j*, PeakVI samples from the area of the latent space that corresponds to *C* and estimates the probability of region *j* being accessible for each sample, then averages over the samples to get a stable estimate of accessibility: 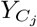 (Methods). Importantly, the representation of the latent space using random variables means that each cell in the original data can be sampled multiple times, allowing PeakVI to sample beyond the available number of observed cells. Additionally, this procedure can be conditioned on batch annotation, thereby correcting batch effects. When comparing two populations of cells, *C*_*A*_ and *C*_*B*_, we use the absolute difference between estimates 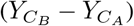 as a measure for the extent of differential accessibility (effect size). Compared to ratio-based statistics (e.g odds-ratio), this estimate is more interpretable (representing absolute increase or decrease in binding propensity) and more stable to low-level signals. For instance, this means that an increase from 0.01 to 0.21 will be equivalent to an increase from 0.7 to 0.9 as opposed to the first being a 20-fold increase and the second being a 1.3-fold increase.

#### 3.1 Using PeakVI estimates for differential accessibility is more sensitive and robust than using the observed data directly

To compare the estimated effect from PeakVI to the empirical effect calculated directly from the observations, we used the hematopoiesis data, and the replicate-batch PeakVI model. We define the empirical accessibility as the proportion of cells in *C* in which *j* is observed as accessible: 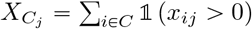, and the empirical effect is defined equivalently to the estimated effect, as 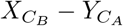. We clustered the latent representations of the cells and ran a series of comparisons for each cluster. First, we ran two comparisons for each cluster: 1) a “biological” comparison, comparing all cells within the cluster to all other cells; 2) an “artifactual” comparison, comparing within each cluster cells that originated from the two large PBMC batches (replicates 1 and 2; excluding clusters with less than 5 cells in either group), (Figure 3A). The biological comparisons are a common use for differential analyses where some real differences in accessibility are expected, whereas the artifactual comparisons are used as negative controls. We ran two additional comparisons for each cluster, comparing cells within that cluster that originated from a given PBMC batch (either replicate 1 or 2) to all cells in all other clusters, which essentially provided two technical replicates of the biological analysis (denoted ‘biological b1‘ and ‘biological b2‘).

**Figure 3:**
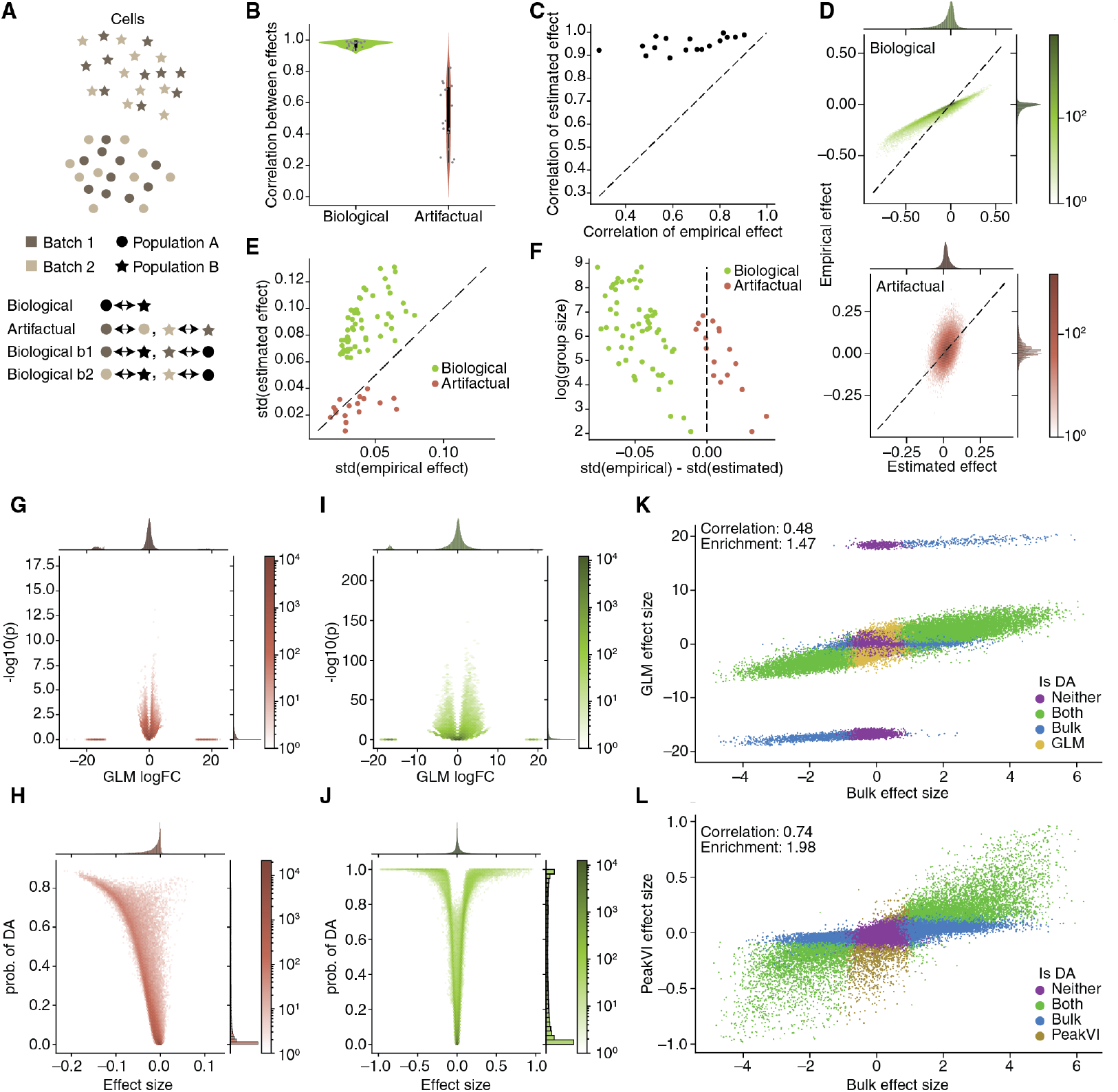
Differential Accessibility Analysis with PeakVI. **(A)** Illustration of the different comparisons. “real”: compare cells between two population; “null”: compare cells from different batches within a single population; “real b1”/“real b2”: compare cells from a specific batch in a population to all cells in the other population. **(B)** Pearson correlations between the estimated and empirical effects. **(C)** correlation of effect size in ‘real b1’ and corresponding effect in ‘real b2’ comparisons. PeakVI estimated effects are far less sensitive to batch effects. **(D)** An example (using cluster 14) relationship between the PeakVI estimated effect to the empirical effect in real (top) and null (bottom) comparisons. **(E)** the width (measured by the standard deviation) of the effect distributions; PeakVI amplifies real differential effects, and silences nuisance ones. **(F)** Level of amplification/silencing depends on level of noise in the empirical effect. **(G-H)** Volcano plots for a GLM (G) and PeakVI (H) when comparing between two batches of NK-cells. **(I-J)** Volcano plots for a GLM (I) and PeakVI (J) when comparing between B-cells and NK-cells. **(K-L)** PeakVI (L) effect is better correlated with a bulk-ATAC based ground truth comparison, and the significant regions have a higher enrichment scores, compared with the GLM (K).

We first measured the correlation between the PeakVI estimated effects and the raw data (empirical) effects. We found that the effects are highly correlated in biological comparisons (mean Pearson correlation 0.97), but less so in artifactual comparisons (mean correlation 0.52) (Figure 3B). We then used the results from “biological b1” and “biological b2” results, and found that the estimated effect is highly reproducible (mean correlation 0.95), while we see a marked decrease in reproducibility of the empirical effect (mean correlation 0.66) (Figure 3C). We also noticed that while the results were highly correlated, there was a difference in the width of the distributions between the estimated and the empirical effects (Figure 3C, S5). To investigate this effect more thoroughly, we calculated the standard deviation of the distributions for each comparison, and found that in all biological comparisons (including “biological b1” and “biological b2”) the estimated effect had a wider distribution than the empirical effect, whereas in artifactual comparisons the distributions were either similarly wide or the estimated effect had a narrower distribution (Figure 3D). We additionally found that this is related to the number of cells included in the compared groups, especially in comparisons that rely on small numbers of cells: in these cases we observed the least difference in standard deviations for the biological comparisons, and the most difference for the artifactual comparisons (Figure 3E).

Taken together, these results demonstrate that PeakVI is amplifying the empirical effect when the effect corresponds to real biological difference, but silences it when it’s a product of noise. When the empirical effect is more susceptible to noise (e.g., smaller number of cells included in the comparison), PeakVI is less able to amplify biological signal, but more efficient in silencing the noise. In contrast, when the empirical effect is calculated with a large number of cells, and is therefore less noisy, PeakVI has less silencing effect, but is able to amplify real differences better.

#### 3.2 Statistical significance with PeakVI

To estimate the statistical significance of differential effects, PeakVI uses techniques described in previous methods from our group [20, 21]. Briefly, during the sampling procedure described above, PeakVI considers pairs of samples, one from each of the compared groups *y*_*a*_, *y*_*b*_. PeakVI determines for each pair if the measured effect for each region *j* is greater than some minimal effect size 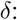 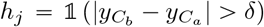 (for one-sided tests: 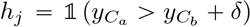). We repeat this many times, and define the probability of differential accessibility, 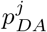, as the proportion of pairs for which *h*_*j*_ = 1 (Methods). We then use a conservative multiple hypothesis correction procedure previously described by Lopez et al. [21] to identify differentially accessible regions with some nominal false discovery rate.

Established pipelines perform this analysis using standard linear models [14] and standard statistical tests like the Wilcoxon rank-sum test or a two-sided T-test [9]. We therefore compared out differential accessibility analysis with a generalized linear model (GLM): a standard logistic regression with an additional covariate for the number of fragments in each cell to avoid library size effects dominating the analysis (Methods). We performed two comparisons using both methods: (1) an artifactual comparison, using the hematopoiesis data we compared between cells from the two PBMC replicates that mapped to cluster 1, corresponding to cells the NK-cell label (Figure 3G-H); (2) a biological comparison, comparing cells from the NK-cell sample to cells from the B-cell sample (using only cells that were FACS-sorted) (Figure 3I-J). We found that both approach show a clear relationship between effect size and statistical significance in both analyses. The GLM results also revealed two common issues, in both comparisons: i) some regions have a very large effect size but are not statistically significant, corresponding to regions that have very low detection rates in both populations; ii) the p-values were inflated due to the large sample size. The GLM model identified 910 regions as differentially accessible in the artifactual comparison, where no biological signal is expected, and 33679 (48.9%) of the regions were differentially accessible in the biological comparison. In contrast, PeakVI results exhibit no numerical issues, and the analyses identified no regions as differentially accessible in the artifactual comparison, and 11362 (16.5%) regions in the biological comparison.

We then ran an equivalent comparison between B-cells and NK-cells using bulk ATAC-seq data from Calderon et al. [22] with sorted immune cell populations, as a ground truth (Methods), and compared the results with the scATAC-seq based results from both analyses (Figure 3I-J). Overall both sets of results are consistent with the bulk results, but PeakVI achieves higher correlation between the effect sizes (0.74 compared with 0.48 for the GLM results), and notably, despite identifying fewer regions as differentially accessibly, 85.8% of the PeakVI results were also differentially accessible in the bulk reference, compared with 65.6% for the GLM. In terms of overlap between the regions found with bulk comparison vs. single cell, both analyses resulted in sets of regions that are over-represented at the bulk results, with PeakVI reaching an odds-ratio of 1.98, and the GLM reaching 1.47. overall, PeakVI provides a well-calibrated statistical significance estimation and enables identification of differentially active regions at a single-region resolution.

### 4 PeakVI unlocks multiple approaches for annotation and discovery of cell states

A major challenge in analyzing scATAC-seq data is the lack of region-based annotations of cell state, in contrast to the abundant resources for RNA-based annotation. Current methods therefore rely on annotations that were generated from gene expression profiles, which are useful but only provide a partial solution, since chromatin accessibility may carry information that is not discernible from gene expression alone. We therefore set out to demonstrate two different approaches for how PeakVI can be leveraged for annotation and downstream discovery. First, PeakVI’s integration capabilities can be used for transfer learning, projecting annotated reference data and un-annotated query data onto a joint space, and transferring insights from the former to the latter. Importantly, this approach relies solely on the regions, without associating regions to target genes or identifying harbored motifs. Secondly, in the lack of an annotated reference, PeakVI’s differential accessibility analysis can be leveraged for de-novo annotation, associating marker regions with nearby genes and identifying enriched gene sets or known marker genes.

PeakVI can be used for transfer learning, by leveraging an annotated reference dataset to annotate a query dataset. First, the reference and query datasets need to be integrated into a joint space, which can be achieved using PeakVI in one of two ways: (i) naively, by analyzing both datasets together and conditioning on the dataset of origin; (ii) using a two-step procedure first presented in scArches[23], in which the reference data is processed in advance, and then incoming query data can be projected onto the reference-based space. The scArches procedure is particularly useful when creating a detailed atlas to be used as a reference resource. After the query and reference are in a shared space, transferring annotations from one to the other can be done using proximity based classifiers, such as KNN or cluster majority vote (which we utilized here). We demonstrate this ability using the hematopoiesis data as the reference, and a dataset of human PBMCs provided by 10X as a query (note that this dataset is different from the multiomic dataset used in previous sections). Notably, the reference data covers both bone marrow and blood, and consists of samples that were sorted to specific cell types, as well as samples that consist of the entire PBMC compartment. We therefore expect the query data to align only to the parts covered by the reference PBMC samples, and not next to cell subsets that are more abundant in the bone marrow. Furthermore, we expect technical hurdles to complicate the integration of the datasets as they were generated by different experimental protocols and processed with different computational pipelines.

We began by creating a reference model, by analyzing the hematopoiesis data using PeakVI in a scArches-compatible configuration (Methods). We then used PeakVI to project the query PBMC data onto the reference space. PeakVI was able to mix the datasets well, only mapping query cells onto areas of the space occupied by PBMCs, but not those corresponding to progenitor cells, which are absent from the query PBMC data (Figures 4A, S6). We then clustered the cells and assigned each cluster with the most abundant cell-specific FACS-based label in that cluster from the reference data. Importantly, these annotations are based on similarity of chromatin landscapes between cells in the query and reference data, without any association to other biological features or aggregation, resulting in a straight-forward labelling of the query data (Figure S7).

**Figure 4:**
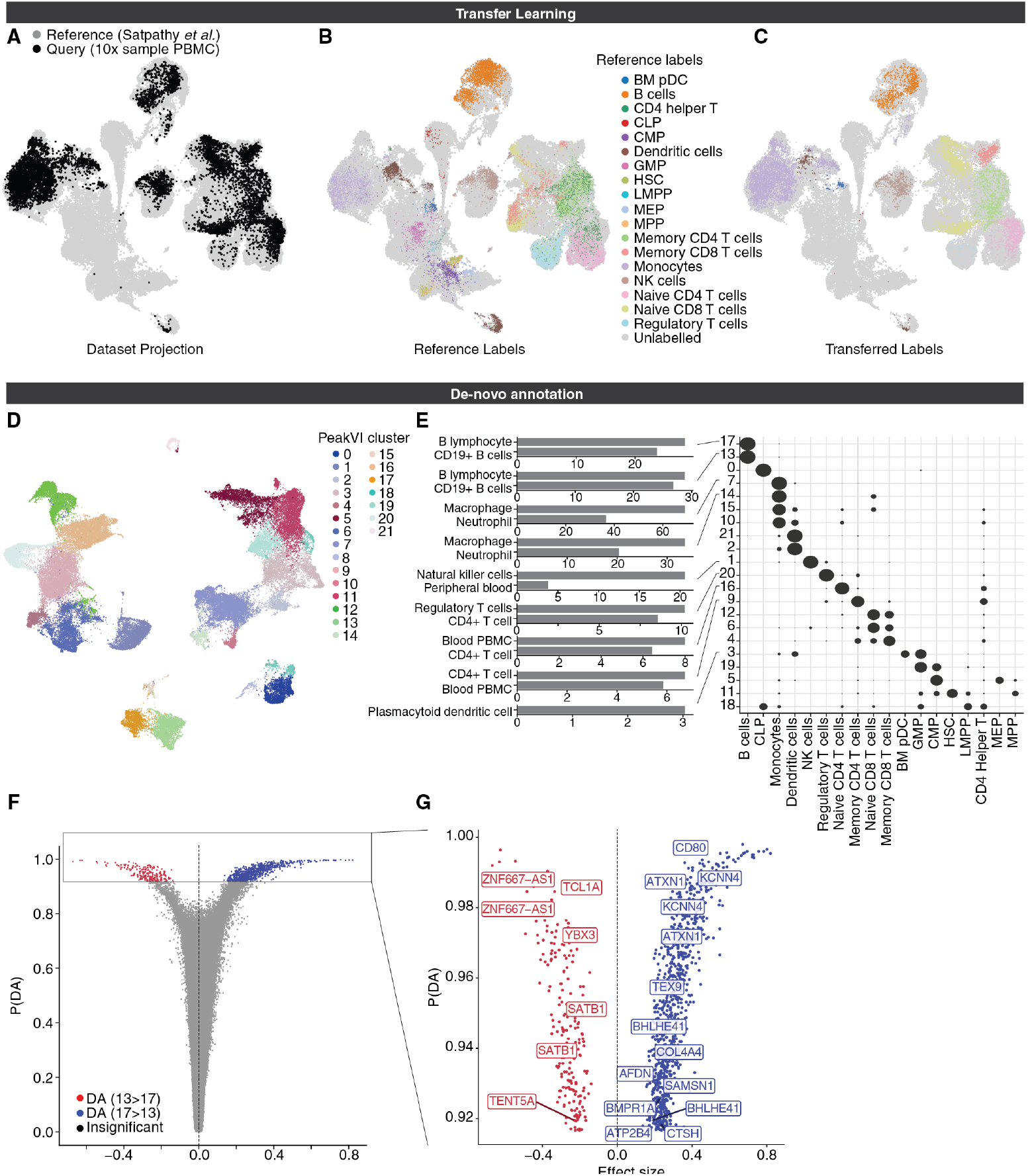
PeakVI unlocks multiple paths for annotation and identification. **(A-C)** PeakVI supports transfer learning. **(A)** Mapping of query data (Sample PBMC data from 10X Genomics) unto a reference data (from Satpathy et al[6]). PeakVI mixes the query data with the reference despite the data being generated by a different protocol and processed by a different pipeline. **(B)** The reference data, colored by FACS-based cell type-specific labels; **(C)** The query data, colored by the transferred cell type-specific labels. **(D-F)** De-novo annotation using PeakVI’s differential accessibility analysis. **(D)** Hematopoiesis data colored by clusters. **(E)** Regions that are preferentially accessible in each cluster were analyzed for enriched cell-type signatures from ARCHS[26] signatures, using enrichr[24, 25]. Heatmap shows distribution of cell type-specific labels for each cluster, normalized by row. **(F)** Volcano plot for a differential accessibility analysis between the two B-cell clusters (clusters 13 and 17). **(G)** Volcano plot for only significant regions, labelled by associated genes that are implicated in Naive B-cells (red) and Memory B-cells (blue).

However, this procedure requires an annotated atlas from a corresponding system, while many scenarios require de-novo annotation, which PeakVI facilitates using the differential accessibility analysis. We demonstrate this using the hematopoiesis data, by de-novo annotating the data and using the FACS-based labels as a ground-truth. We first clustered the latent space (Figure 4D), and consistent with our previous findings we found that clusters tend to consist primarily of cells that have the same label. Next, using our differential accessibility analysis, we compared each cluster to all other clusters except for the 3 most similar clusters, to avoid highly-similar clusters masking the differences (Methods). For each cluster we used a one-sided test to only identify regions that are preferentially open in the target cluster. We then used enrichr [24, 25] to associate the regions to nearby genes and leveraged the ARCHS4 [26] collection to find over-represented cell-type specific gene signatures. We were able to confidently identify many of the cell type-specific clusters, which matched their FACS-based label (Figure4E, Methods). For instance, marker regions for clusters 13 and 17, in which labelled cells are overwhelmingly B-cells, were indeed enriched for regions associated with B-cell marker genes; Cluster 1 marker regions were enriched for NK-cell marker genes, and indeed the labelled cells in that cluster are NK-cells. Similarly signatures for CD4+ T-cells, Regulatory T-cells, and pDCs, were all highly enriched in the clusters with the corresponding FACS-based labels. Thus, using PeakVI and gene-based signatures, we are able to annotate the data and recapitulate many of the FACS-based labels.

These results are nonetheless limited by the availability of gene signatures, which may not be available for all cell types, or provide only a a high-level annotation at a limited resolution. Specifically, most progenitor cells in the hematopoiesis data could not be annotated in a similar fashion for lack of corresponding signatures, and despite clustering separately, both CD4+ naïve T-cells and CD4+ memory T-cells were annotated simply as CD4+ T-cells, since higher-resolution signatures were not available. PeakVI can therefore be used in a two-step approaches whereby cells can be stratified into broad types, using reference-based annotation, and then assigned with more high resolution labels of cell sub-types or states using de-novo analysis. As a case in point, we focused on the set of cells which were annotated as B cells in our reference-based analysis. These cells can be divided into two clusters (clusters 13 and 17). To derive a higher resolution annotation of the B cell compartment, we ran a two-sided comparison between the two clusters and identified 1043 differentially accessible regions in total, 207 preferentially accessible in cluster 13 and 836 preferentially accessible in cluster 17 (Figure 4F; Supp Table 2 and Methods). Among the genes associated with regions detected for cluster 13 we found TCL1A, known to be expressed throughout B-cell differentiation up to naive B-cells but silenced in memory B-cells and plasma cells [27, 28], and YBX3, implicated in B-cell differentiation as an immature B cell marker [29]. We also found SATB1, TENT5A, and ZNF667-AS1, which along with TCL1A and YBX3, were previously found to be differentially expressed in naive B-cells compared with Memory B-cells [30]. Concordantly, genes associated with cluster 17 included known markers for memory B-cells AIM2[31] and CD80 [32], and 9 other genes previously found to be differentially expressed in memory B-cells compared with naive B-cells [30] (Figure 4G). Taken together, we concluded that cluster 13 consists of naive B-cells and cluster 17 consists of memory B-cells, therefore demonstrating that PeakVI’s differential accessibility analysis can be used in conjunction with a reference-based annotation to increase the resolution of annotations and identify new targets for further study.

## Discussion

PeakVI is a deep generative model for analyzing single cell chromatin accessibility data. The model is designed to explicitly account for various technical effects that mask and distort the biological signal. The latent representation learned by the model is probabilistic in nature, embedding the observed cells in a smooth variational space that preserves the biological heterogeneity, minimizes confounding effects, and can be used directly to explore the chromatin landscape of a population of cells.

PeakVI improves upon previous attempts to use deep learning to analyze scATAC-seq data in several manners. First, the architecture used in the underlying neural networks scales with the size of the input data, increasing the expressiveness of the model to match with increasingly large and complex datasets (Methods). Second, PeakVI accounts for technical confounders and enables correction of batch effects, with clear benefits to downstream results. Thirdly, Since it is common for features (regions) to outnumber the samples (cells), and the observations are mostly binary and therefore contain little information, PeakVI also takes measures to successfully prevent the model from over-fitting, by holding out some of the data as a validation set, tracking the model’s performance on the validation data, and halting the training process when the performance on the validation data stops improving, thus ensuring that the model is learning generalizable features. Finally, PeakVI provides extensive methods to take advantage of the learned latent space for analysis tasks beyond dimensionality reduction, visualization, and clustering. Specifically, PeakVI enables high resolution annotation of cell state, by allowing both reference-based analysis and de-novo annotation analysis. In that capacity, PeakVI enables accurate differential accessibility analysis at a single-region resolution that reduces the effect of confounders and avoids common issues with the current practices for differential accessibility, namely numerical instability and inflation of significance scores.

PeakVI is robust to low-quality data, easy to configure, train, and use. It is implemented in the scvi-tools suite[15], which provides interfaces with popular processing environments like scanpy[33] and Seurat[34]. As such, PeakVI can be easily incorporated in existing analysis pipelines to enhance current analyses for dimensionality reduction, batch correction, differential accessibility, and annotation.

## Declarations

### Acknowledgements

We thank Florian Wimmers for many helpful discussions and insightful feedback. We thank Christina Usher for assistance with visualizations.

### Author contributions

TA and NY conceived of the model and designed the analyses. TA implemented the model with input from AG. TA and DAR performed the analyses. NY supervised the work. TA and NY wrote the manuscript.

### Data and Software Availability

All data used in this manuscript is publicly available via the original publications and releases. Intermediate data, trained models used in this manuscript, and the notebooks to generate the figures in this manuscript, are all posted and available on zenodo: 10.5281/zenodo.4728534.

### Conflicts of interests

All authors declare that they have no competing interests.

## Methods

### 5 The PeakVI Model

Let 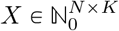 be a scATAC-seq region-by-cell matrix with *N* cells and *K* regions, where *x*_*ij*_ ∈ ℕ_0_ is the number of fragments from cell *i* that map to region *j*. Since PeakVI models the probability of observing a region, regardless of the number of reads supporting that observation, the observations are treated as binary: *X*\* ∈ [0, 1]^*N* × *K*^, where 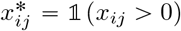. The observations are therefore generated from a Bernoulli distribution *x*\* ~ *Ber(q*_*ij*_). PeakVI computes *q*_*ij*_ as a product of three probabilities: *q*_*ij*_ = *y*_*ij*_ · *r*_*j*_ · *ℓ*_*i*_, where *y*_*ij*_ captures the true biological heterogeneity; *r*_*j*_ captures region-specific biases (e.g width, sequence); *ℓ*_*i*_ captures cell-specific biases (e.g library size). The three probabilities are estimated jointly using deep neural networks.

The biological component *y*_*ij*_ is estimated using a VAE[17], which is composed of two deep neural networks, the encoder *f*_*z*_ and decoder *g*_*z*_. Briefly, the encoder 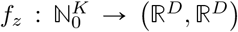, computes the distributional parameters of a D-dimensional multivariate normal random variable: *Z* ~ *MVN*(*f*_*z*_(*x*_*i*_)_1_, *f*_*z*_(*x*_*i*_)_2_). The sample is then concatenated to the batch annotation for cell *i*, and passed through the decoder *g*_*z*_ : (ℝ^*D*^, {0, 1}^*S*^) → [0, 1]^*K*^, for *S* being the dimension of the one-hot batch annotation (the number of batches). The cell-specific factor *ℓ*_*i*_ computed from the input data for cell *i* via a deep neural network 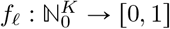. Finally, the region-specific factor *r*_*j*_, since it is optimized across samples, is stored as a *K*-dimensional tensor, used and optimized directly.

#### 5.1 Architecture

All PeakVI neural nets are fully connected networks, composed of repeated blocks that share a basic structure. For convenience, we define a fully connected block *FC(I, O, D, A)* as having a fully connected layer with *I* input nodes and *O* output nodes, followed by a drop-out layer with a *D* probability of dropout, a layer-norm layer, and finally an *A* activation function.

The encoder *f*_*z*_ is constructed as follows:

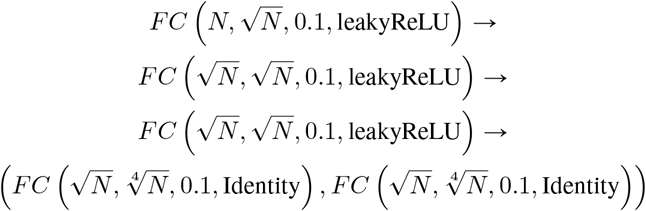

With 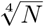 being the default dimensionality of the latent representation. This ensures that the model architecture scales with the number of features in the data and the complexity of the representation.

The decoder *g*_*z*_ is constructed as follows:

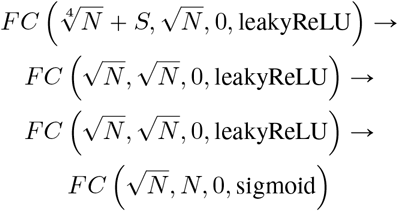

With *S* as the dimensionality of the batch annotations, concatenated to the latent representation.

The cell-specific factor network *f*_*ℓ*_ is constructed similarly:

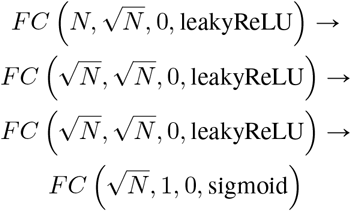

#### 5.2 Training Procedure

By default, PeakVI is optimized using AdamW[35] with a learning rate of 0.0001, weight decay of 0.001, and minibatch size of 128. The model is trained on 90% of the data, with the remaining 10% used as a validation set. Training is performed for at most 500 epochs, with early stopping: if there is no improvement in terms of the reconstruction loss on the validation set for 50 epochs, the training stops. For epochs *i* ∈ [1, 50] the KL divergence term is weighed done by a factor of *i*/50. The best state throughout training, defined as the state that achieves the best reconstruction loss, is saved during the training and used as the final state. All training settings are configurable.

#### 5.3 Differential Accessibility Analysis

For a differential accessibility analysis between two populations *A* and *B*, the analysis is performed as follows:

1. *N* cells are sampled from each population, with replacement (default *N* = 5000). We denote the resulting cells 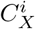 for the *i*-th sample from population *X*, for *i* ∈ [*N*] and *X* ∈ {*A, B*}.
2. for each cell *C*, we apply the inference model on the cell’s chromatin accessibility profile *f*_*z*_(*x*_*C*_) to get the variation distribution corresponding to that cell, *q*_*C*_, sample from that distribution to get an estimated profile of the probability of accessibility of all regions in that cell: *z*_*C*_. We then use the generative model *g*_*z*_ to estimate the probability of accessibility of each region *j* in that cell: (*yC*)_*j*_. Sampling from the variational space allows us to sample the same cell multiple times and get different estimates, thereby enabling statistical power beyond the original sample size.
3. to calculate the effect size for each region, we simply take the average estimated probability of accessibility across all samples from each population, and compute the absolute difference between the averages: 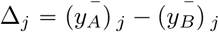.
4. to calculate the statistical significance, we randomly pair samples from each population into *N* pairs of estimates {(*y*_*A*_, *y*_*B*_)^*i*^ | *i* ∈ [*N*]}, then for each region we count for how many pairs the difference between estimates was greater than some minimal *δ* (default 0.05): 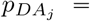 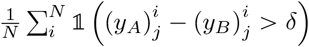. This procedure has been previously described by [21].
5. In addition to *p*_*DA*_, we also compute the Bayes factor: 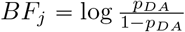, and perform multiple testing correction using the procedure previously described by Lopez et al[21] to get a qualitative, binary label for each region.

### 6 Benchmarking and Evaluation

#### 6.1 Stability Analysis

To measure the stability of PeakVI to hyperparameter selection, we ran a full grid search using the 10X Genomics sample data. We held out 10% of the data as a test set and trained all models on the remaining set. We trained each model 3 times (with an independent train-validation split) and measured the likelihood on the held-out data. The full results are available in Supplemental Table 1. The hyperparameters we varied and the values used are as follows: learning rate (1e-2, 1e-3, 1e-4); number of hidden layers (1,2,3,4); dropout rate (0.1, 0.3); minibatch size (64, 128, 256); weight decay (0.1, 0.01, 0.001).

#### 6.2 Dataset Processing

The hematopoiesis data was downloaded from GEO (Accession GSE129785); specifically the processed peak-by-cell matrix and metadata files: scATAC-Hematopoiesis-All.cell-barcodes.txt.gz, scATAC-Hematopoiesis-All.mtx.gz, scATAC-Hematopoiesis-All.peaks.txt.gz. We then filtered the genomic region to only those that are detected in at least 0.1% of the cells in the sample, reducing the data from 571400 regions to 133962 regions. The sample data from 10X genomics was also downloaded as preprocessed peak-by-cell matrices, without any additional filters.

#### 6.3 Running Published Methods

For all methods, we followed the standard recommended procedure for analyzing data. For visualization, we computed the umap[36] coordinates using the python implementation from the latent space computed by the respective method (except for SCALE, see below). **cisTopic**(v0.3.0): We used the WarpLDA model fitting procedure, and chose the best number of topics based on the second derivative, as recommended by the package documentation. For the hematopoiesis data the model used 100 topics, and 40 topics for the paired PBMC sample data from 10X Genomics. **chromVAR** (v1.12.0): We used the JASPAR2016 motif set, containing 386 motifs, and followed the standard analysis outlined in the package documentation. We used the unnormalized motif deviation scores. For dimensionality reduction, we found no clear difference between using the chromVAR scores directly and applying an additional linear procedure (i.e principle component analysis). Results described in the manuscript use the deviation scores directly. **LSA:** We used the python implementation from the Scikit-learn[37]. We first binarized the data, then computed the top 50 components used the TruncatedSVD method, on the tfidf-transformed data. **SCALE** (v1.0.4): we used the external script to run SCALE without a pre-determined number of clusters, using the default arguments. In all visualizations, we used the umap coordinates computed by SCALE.

#### 6.4 Enrichment Score Calculation

Enrichment scores used to quantify cell type separation and batch mixing were computed in an identical way. Given a latent representation *R*, an integer *k*, and cell labels *L*, we first compute *G*_*R,k*_, the K-nearest neighbor graph from *R* with *k* neighbors. We then compute for each cell the proportion of neighbors that share the same label: 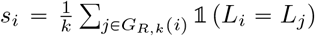. The overall score is the average score across all cells, 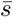, normalized by the expected score for a random sample from the distribution of labels: 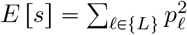, for {*L*} being the set of available labels, and *p*_*ℓ*_ being the proportion of each label *ℓ* ∈ {*L*}. The enrichment score is then 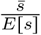.

#### 6.5 Differential Expression with Logistic Regression

As a simple benchmark for differential accessibility, we constructed a standard logistic regression model to compare B-cells to NK-cells, using the design *y* ~ number of fragments + cell type, where *y* is the binary detection of a genomic region. We fit the model using the *glm* function in *R*. Due to the runtime of this analysis, we limited the results to regions that are detected in at least 1% of the compared cells.

#### 6.6 Analysis of bulk ATAC-seq data

The bulk ATAC-seq data used as a ground truth reference for differential accessibility analysis was downloaded from GEO (accession GSE118189). We used the unstimulated samples of all B-cell and NK-cell subtypes included in the study and used DESeq2[38], which was found to be among the ebst performing methods for differential accessibility from bulk ATAC-seq data[39] for differential accessibility between the two group. We then found regions in the hematopoiesis data that overlap with the regions in the bulk data, and used the differential signal found in the bulk data for the overlapping regions in the hematopoiesis data.

#### 6.7 Projection of query data onto reference

Projection of query data onto a latent space learned from reference data is done using scArches[23]. First, the 10X sample PBMC data was downloaded and processed (using CellRanger v3.1.0) using the hematopoiesis peaks. We then trained a PeakVI model on the hematopoiesis data using ell covariate injection, which adds one-hot encoded batch annotation to each layer in the VAE (as opposed to only the decoder layers, which is the default behavior). We then trained the resulting model on the query data, which involves adding batch annotations corresponding to the query data, and only training the nodes in the network that interact with these additional batches. This preserves the latent representation of the reference data while projecting the query data onto the same space, while correcting batch effects between the query and data.

#### 6.8 Cluster Annotation with differential accessibility

Differential accessibility to identify marker regions for each cluster was performed between each cluster and all other clusters except the three most similar clusters. This was in order to avoid sampling pairs of cells that are highly similar from the two groups, which would reduce the signal. We therefore calculated the centroid of each cluster (the average position in the latent space of all cells in the cluster), computed the Euclidean distance matrix between all centroids, and identified for each cluster the 3 most similar clusters. We then used the identified regions (using the Bayesian FDR method described by Lopez et al. [21]), ran them through enrichr[24, 25], and downloaded the enrichment results for the ARCHS4 Tissues set. For associating regions with genes, we used the bioconductor package TxDb.Hsapiens.UCSC.hg19.knownGene[40] and considered only strict overlaps between the region and the annotated gene body or promoter.

---

**Figure S1:**
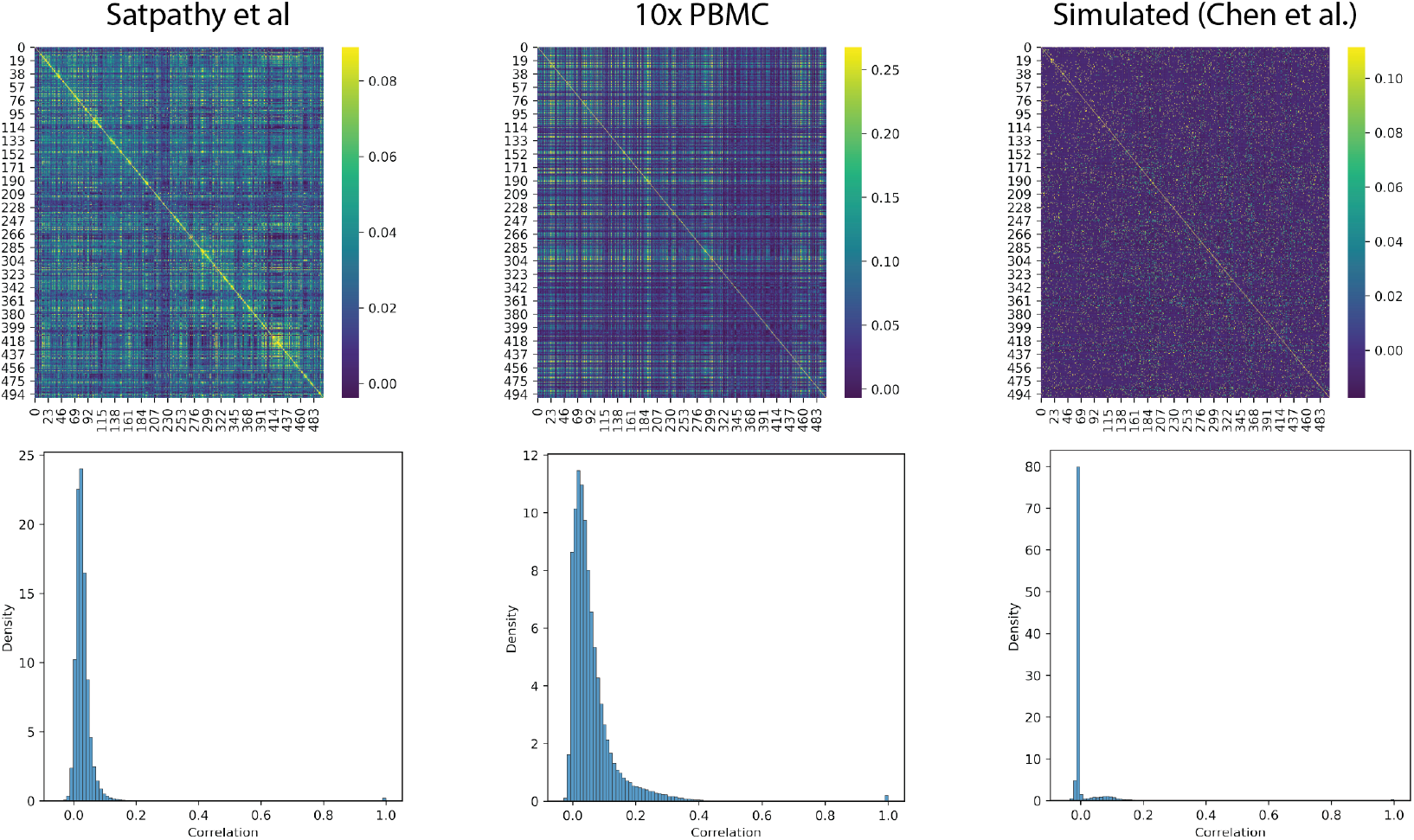
A Pearson correlation matrix (top) and distribution of correlation coefficients (bottom) of regions in three datasets: the immune cell dataset from Satpathy et al [6] (left); the sample multi-omics 10K cells PBMC dataset from 10x Genomics (center); and a simulated Bone Marrow dataset generated by Chen et al [12]. [cite]. For visual purposes, figures were generated using only the first 500 regions in each dataset, and across all available cells. Simulated data does not adequately represent the covariance structure of real scATAC-seq data.

**Figure S2:**
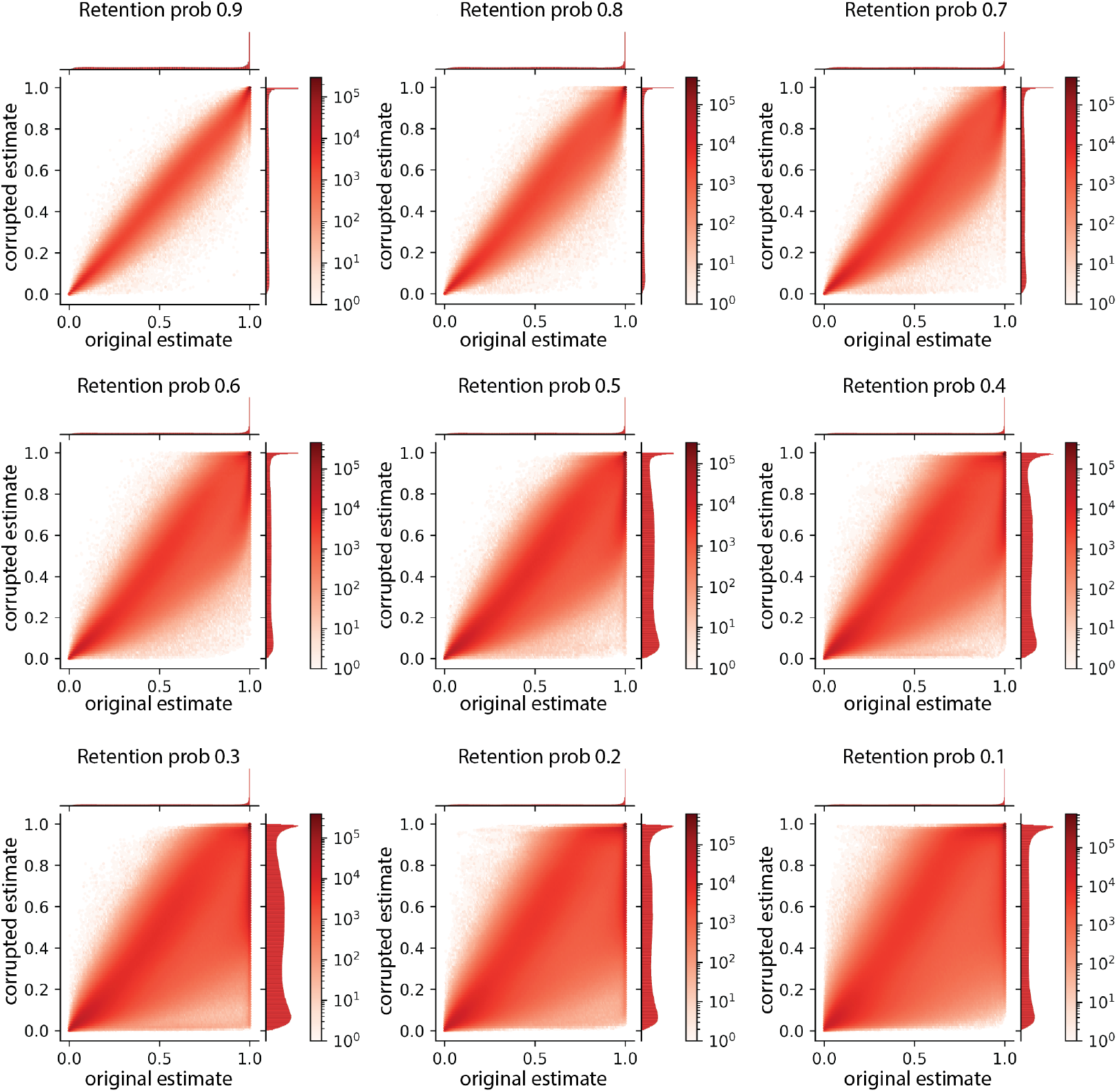
Corruption analysis, in which observations were randomly replaced by zeros. Visualization is limited only to corrupted indices, showing that while increased corruption destabilizes the model, PeakVI is overall highly robust to the sparsity of low quality data.

**Figure S3:**
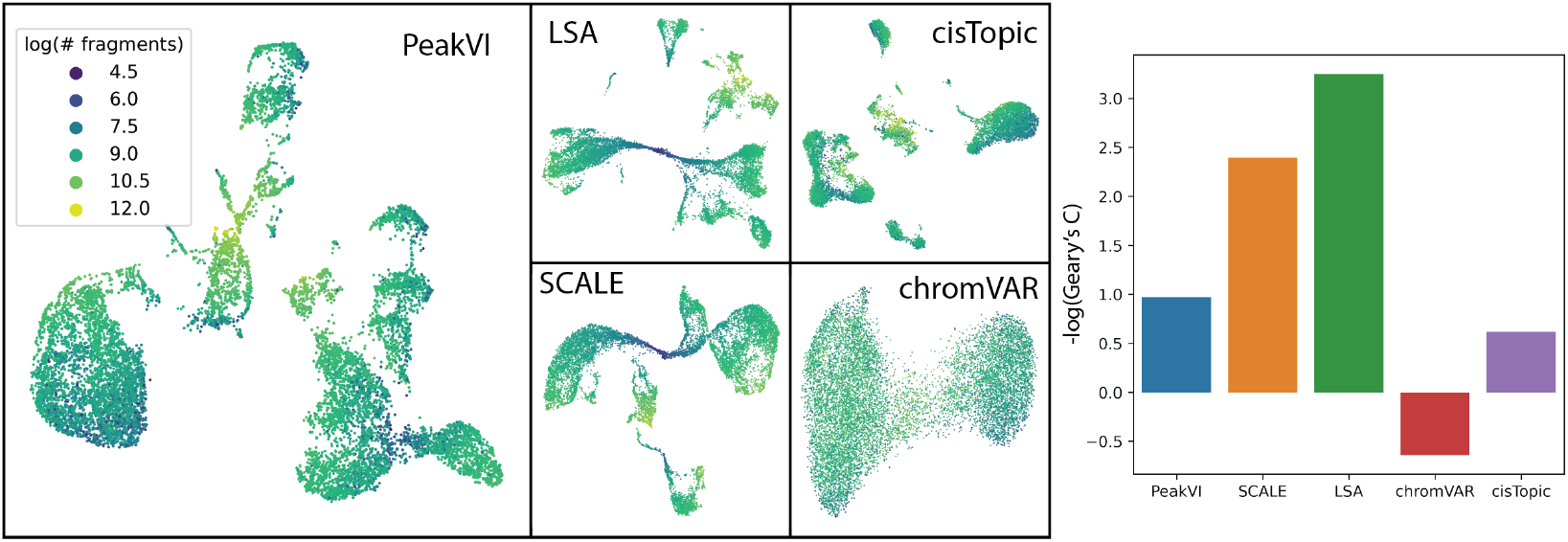
UMAPs of the sample paired scRNA and scATAC-seq PBMC data from 10X genomics, colored by the number of fragments mapped for each cell (left) and the spatial autocorrelation measured using Geary’s C[19] (right). LSA and SCALE are most impacted by library size effects, PeakVI and cisTopic are robust, and chromVAR is negatively correlated.

**Figure S4:**
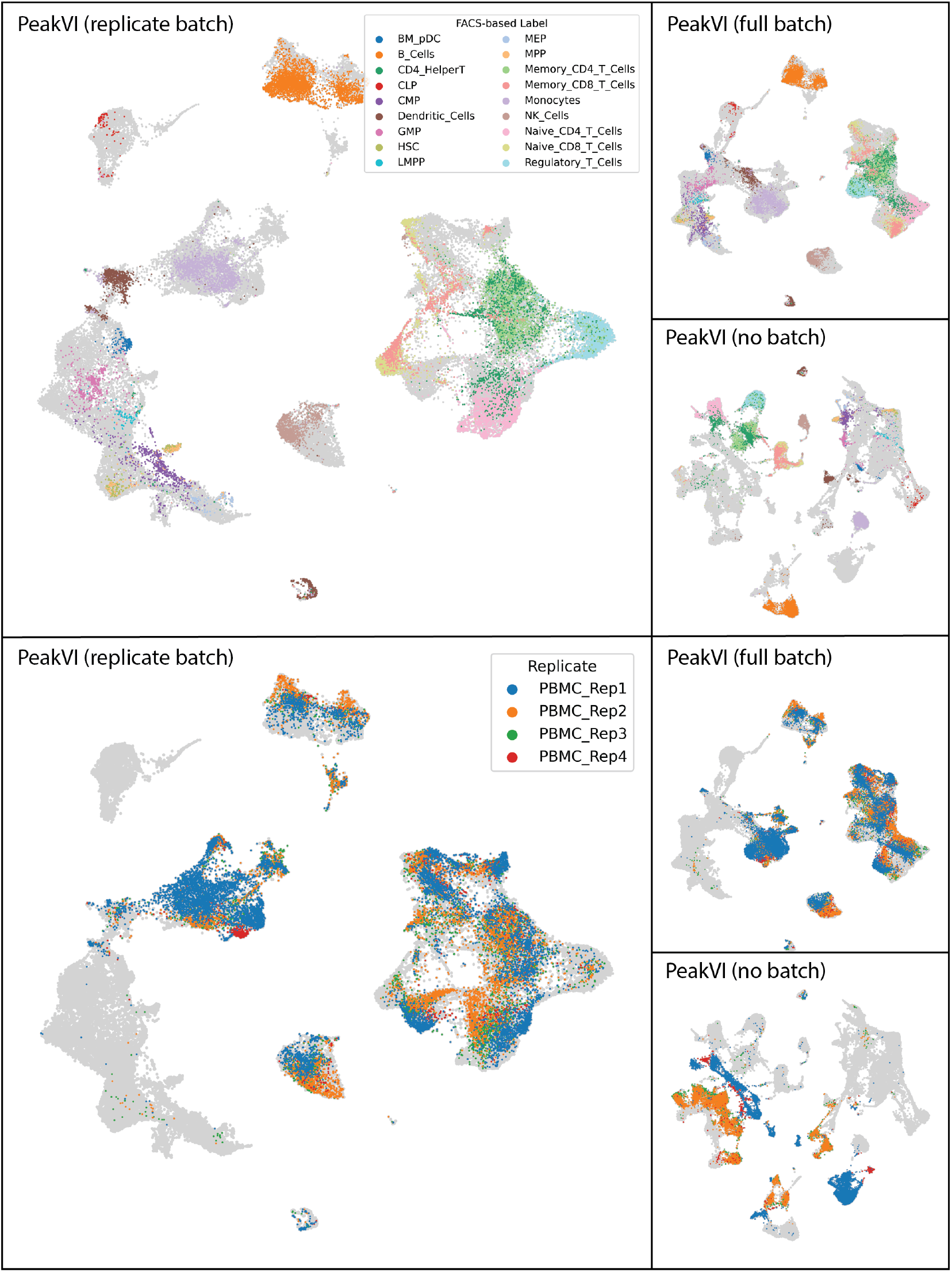
Visualizations of the Satpathy data using three configurations of PeakVI: treating replicates of multi-replicate samples as separate batches (replcate batch); without batch correction (no batch); treating each sample as a separate batch (full batch). Colored by FACS-based labels (top) and replicates of the unsorted PBMC samples (bottom). Unlabelled cells are colored in light gray.

**Figure S5:**
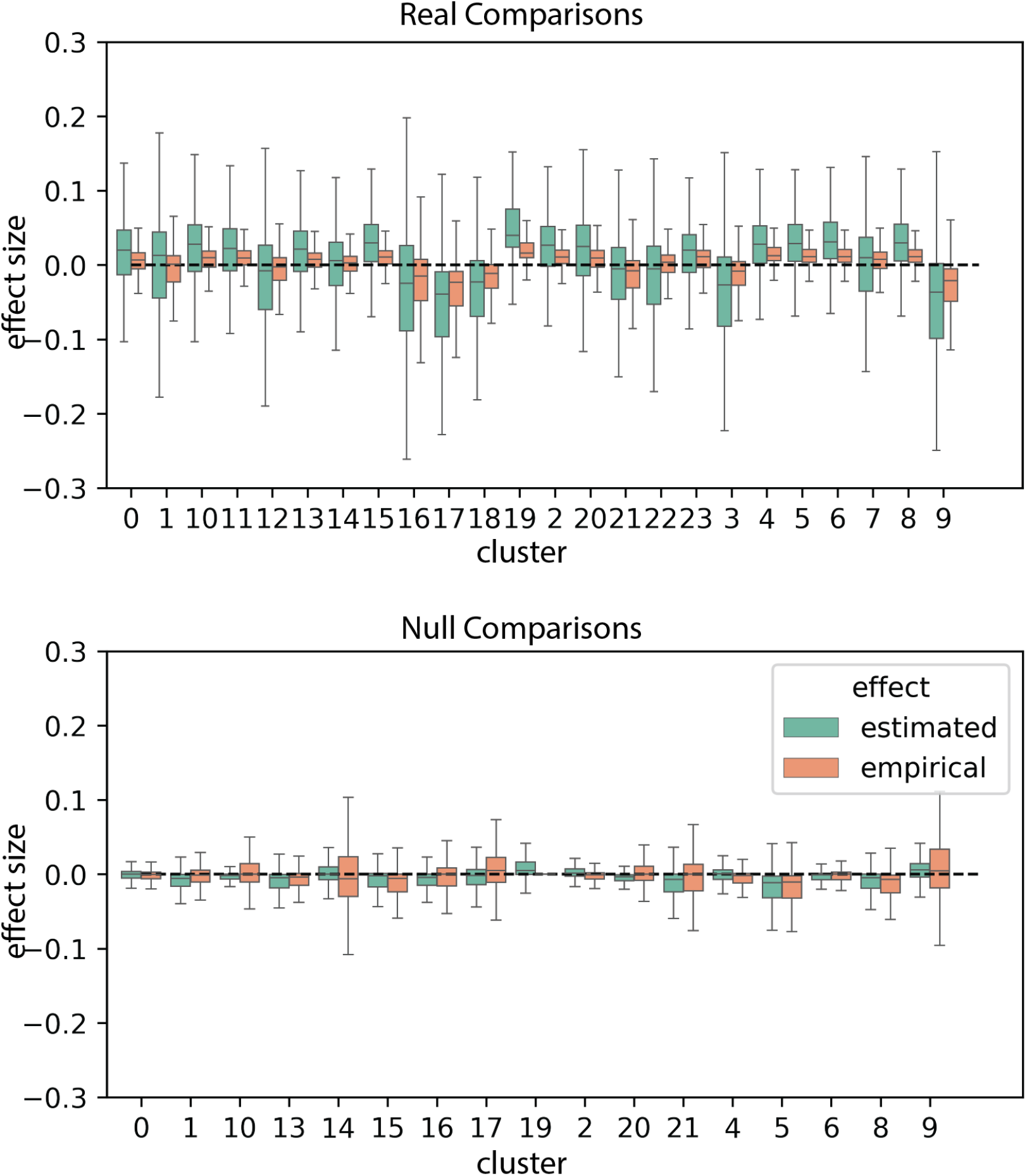
The effect size distribution for each real (top) and null (bottom) comparison. PeakVI estimated effects are amplified compared with the empirical effect in real comparisons, but the opposite is true for null comparisons. Overall PeakVI consistently has a better signal-to-noise ratio.

**Figure S6:**
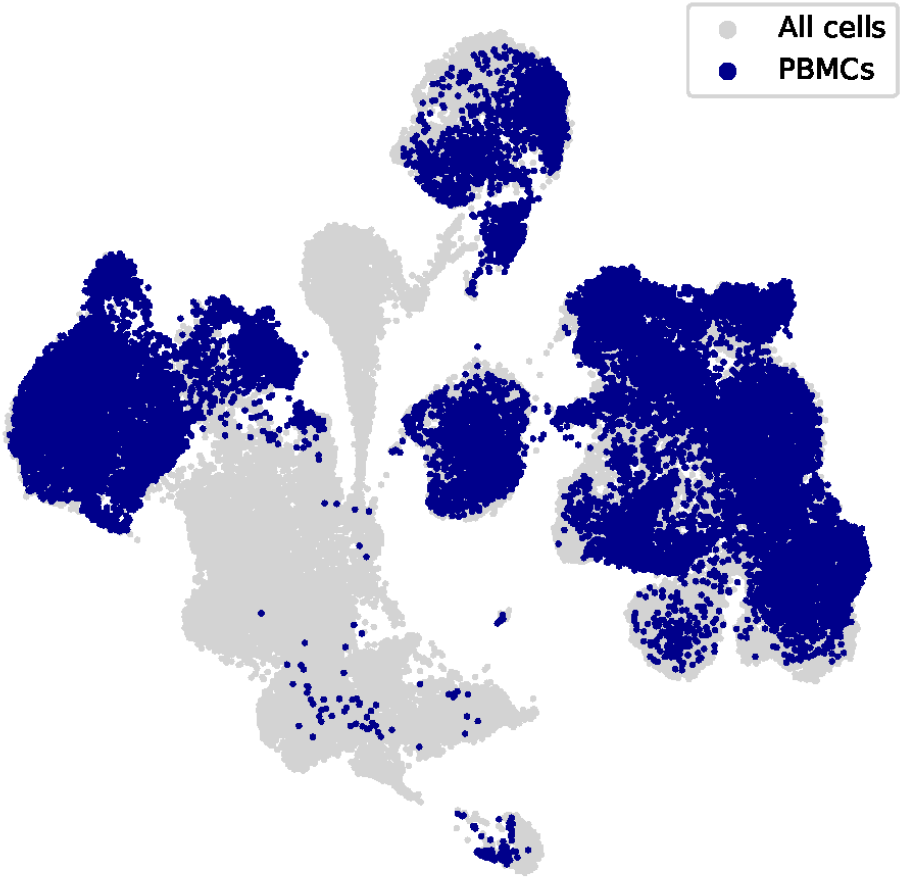
The low-dimensional representation of the Hematopoiesis data, trained in a scArches-compatible manner, with cells from PBMC samples in dark blue, showing how PBMCs are distributed in the space.

**Figure S7:**
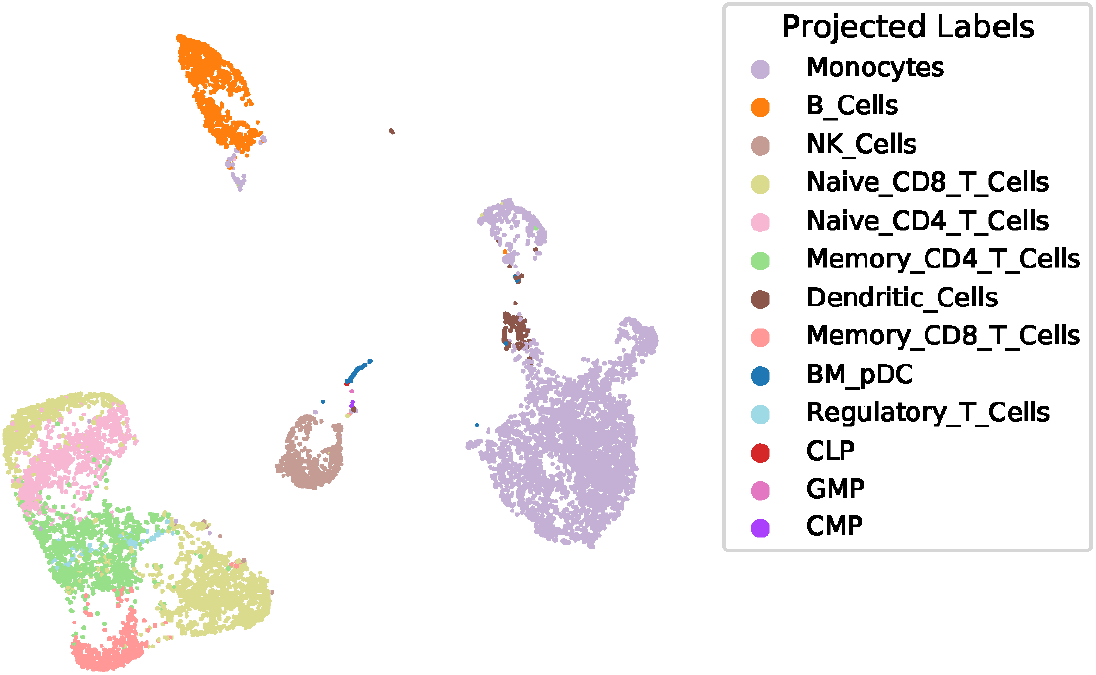
The low-dimensional representation of the Sample 10X PBMC data, with labels transferred from the hematopoiesis data.

## Notes

### Competing Interest Statement

The authors have declared no competing interest.

